# Cascading Effects of Shrimp Trawling: Increased Benthic Biomass and Increase in Net Primary Production

**DOI:** 10.1101/298323

**Authors:** Luczkovich Joseph John, Deehr Rebecca A, Kevin J Hart, Lisa M Clough, Jeffrey C Johnson

## Abstract

Trawling has been shown to cause high mortality of discarded species (bycatch) and short-term ecological disturbance to bottom communities in coastal systems, resulting in lowered benthic biomass. Here we report evidence of a trawling-induced trophic cascade resulting in an increase in biomass of benthic polychaetes after the end of the shrimp trawling season in areas open to trawling in North Carolina (USA). Using comparative measurements of abundance of bycatch species and benthos in open and closed trawling management areas and Ecopath network modeling, we show that trawling in the open area has led to increases in deposit-feeding polychaetes and decreases in bycatch species (fish and crabs) that are benthic predators on the polychaetes. We conclude that proposed management actions to reduce the shrimp trawl fishery effort will influence other net and trap fisheries for southern flounder and blue crabs indirectly, as revealed by our network models, and the proposed trawling ban may lead to improvements in other valuable fisheries.

## Introduction

Globally, bottom trawling is one of the most important types of fishing gear, accounting for 17 % of the global catch of all species in 1990; this is especially true for wild shrimp and prawn fisheries globally, with trawls accounting for 87% of the harvest (1). Trawls have been used since the late 1800’s, first on sailing vessels, then on steam-powered vessels, then on motorized vessels with an increasing amount of fishing power or effort (2). Bottom trawling has been shown to alter marine ecosystems by reducing the abundance of trawled species and disturbing bottom habitats (2,3). Previous studies (4–9) suggest that the benthos should be reduced by action of the trawls, which dig into and ride over the bottom, causing a physical bottom disturbance. For example, a meta-analysis of 59 experimental and observational studies (4) revealed that otter trawling should produce on average a 31% decrease in benthic population densities after a single short-term trawling disturbance event, with this effect being habitat-dependent and somewhat larger (57%) in muddy bottom habitats or smaller (21%) in sandy habitats. That same study revealed variation in the response of taxa of benthos to short-term trawling disturbance effects: corals and crustaceans showed the largest (75 %) declines, whereas polychaetes, ophiuroids, holothurians, echinoids, and gastropods showed intermediate (50-75%) declines, bivalves and sponges showed 40% declines, and asteroids and oligochaetes were most resistant to trawling and showed the smallest declines (20-30%); all species examined showed declines, and none showed increases relative to the controls. In contrast with this meta-analysis, here we report an increase in benthic biomass in areas open to shrimp trawling in a North Carolina estuary when compared with no-trawling areas and in open areas after the trawling season has largely ended. In addition, an ecosystem model was developed for examining the effects of shrimp trawling, which simulated this increase in benthos biomass. The model further indicated an increase net primary production, suggesting a previously unrecognized beneficial effect of bottom trawling.

The trawling impacts on the benthos are also expected to cause or can be caused by fishing-induced trophic cascades. Trophic cascades are indirect community and ecosystem impacts that occur due to dramatic changes in abundance of a species at one trophic level (i.e., predator removal) that affect species at two or more trophic levels. Fishing practices, such as trawling, seining, spearing, and even recreational fishing, have been implicated in causing trophic cascades in marine ecosystems (10–16). However, trawling in soft-bottom ecosystems such as North Carolina (USA) estuaries has not been demonstrated previously to cause a trophic cascade. Trawling for shrimp is widely practiced in the southeastern USA and elsewhere (17–21) and has a large potential impacts from sediment re-suspension, removal of bycatch and associated discards, and has great potential to cause trophic cascades.

The questions we ask here are what happens to an entire estuarine ecosystem where trawling has taken place over many years? Does the benthos show declines as previously described in short-term trawling experiments? How do the populations of higher trophic level predators respond? Is there a trophic cascade that occurs when shrimp trawling on soft-bottoms is practiced repeatedly? And can we simulate and verify these dynamic processes? We report a 200% measured increase in benthos in a heavily trawled area at the end of the shrimp trawling season when it is compared to nearby, otherwise similar, areas closed to shrimp trawling in terms of the density and biomass of benthic polychaete worms. Furthermore, small fish and crab benthos-feeding predators, commonly caught in shrimp trawls and discarded as bycatch, were in lower abundance in trawled areas relative to untrawled areas after the trawling season, which implies that trawls can act like large predator on the by-catch species, removing them, and initiating a trophic cascade. Finally, we used ecosystem trophic network visualizations and simulation models to show that this increase in benthos after trawling is likely to be due to the cumulative effects of a trawling-induced trophic cascade, due to the removal of predators during trawling and a scavenger subsidy effect due to the discarded bycatch from trawling, which feeds the benthos and crabs. This new result shows that discards and trawling disturbance may have different long-term effects than short-term trawling experiments have shown at the whole-system level.

## Methods

### Ecopath Modelling Procedures

The ecological network models were built in Ecopath with EcoSim v 6.4 using data that we collected on various species across the trophic spectrum and group biomasses and those biomasses estimated from commercial harvest data. Commercial harvest data were obtained from the North Carolina Division of Marine Fisheries (NCDMF) Trip-Ticket Program database for 2006-2008, including all commercial species harvested in the Core Sound Management Area. The NCDMF trip ticket is a form used by fish dealers to report commercial landings information. Trip tickets collect information about the fisherman, the dealer purchasing the product, the transaction date, the number of crew, area fished, gear used and the quantity of each species landed for each trip.

Ecopath network models were built for the Core Sound Management Area, using data from 2007 measured in the spring and fall seasons (before and after the shrimp trawling peak in July) for areas open to trawling and during the spring and fall seasons for closed trawling areas, a total of four seasonal models. Two additional annualized models were created, with identical compartments, one for areas open to trawling and one for areas closed to trawling. Each model had 63 living compartments with the biomasses (in g C m^−2^) of various species (with some compartments comprised of aggregated species groupings) and two compartments with non-living carbon (bycatch and detritus) (Table 2). Bycatch data were obtained from measurements taken as part of an observer program in the shrimp fishery of Core Sound (22). Detritus was directly measured (see below).

**Table 1.**
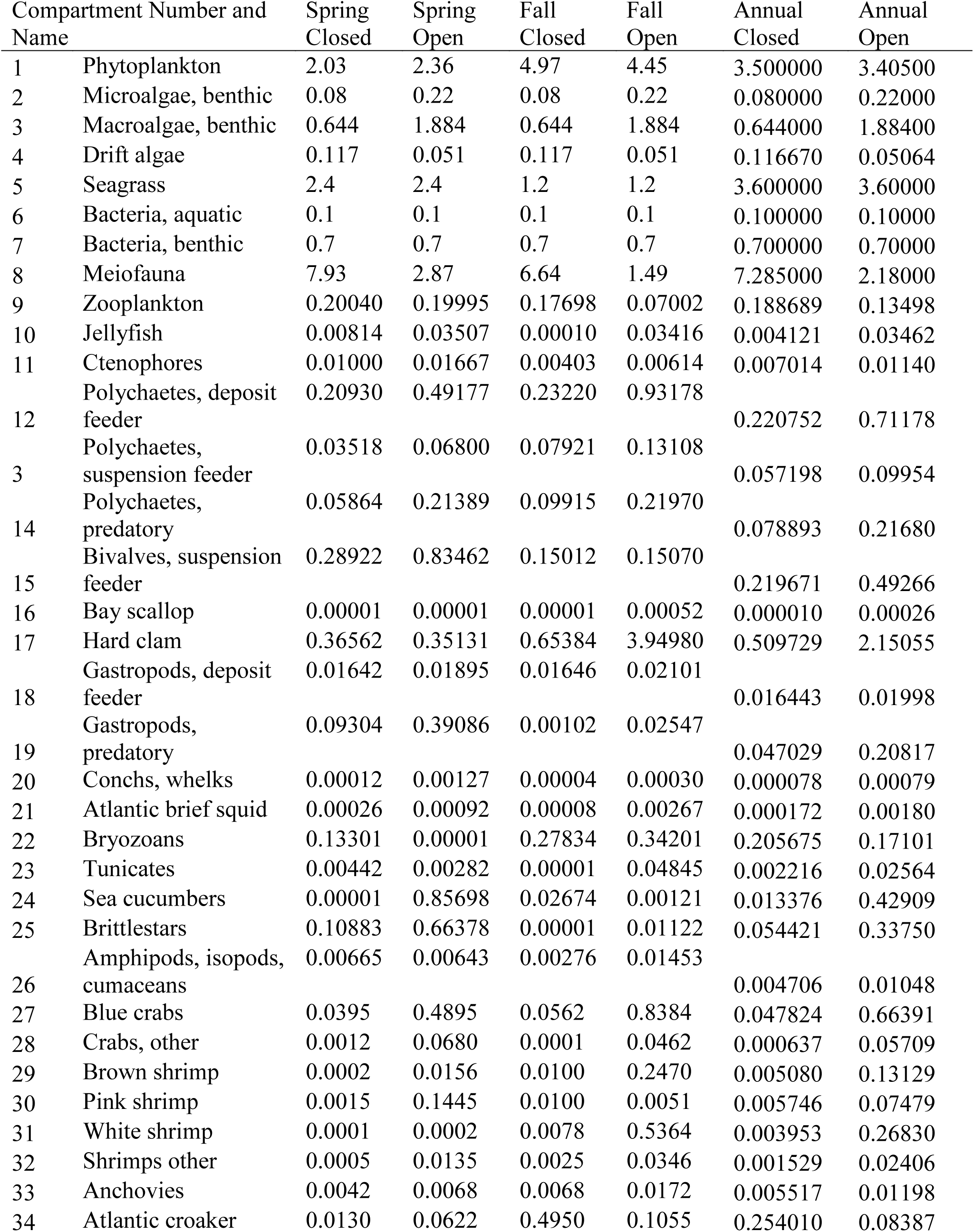
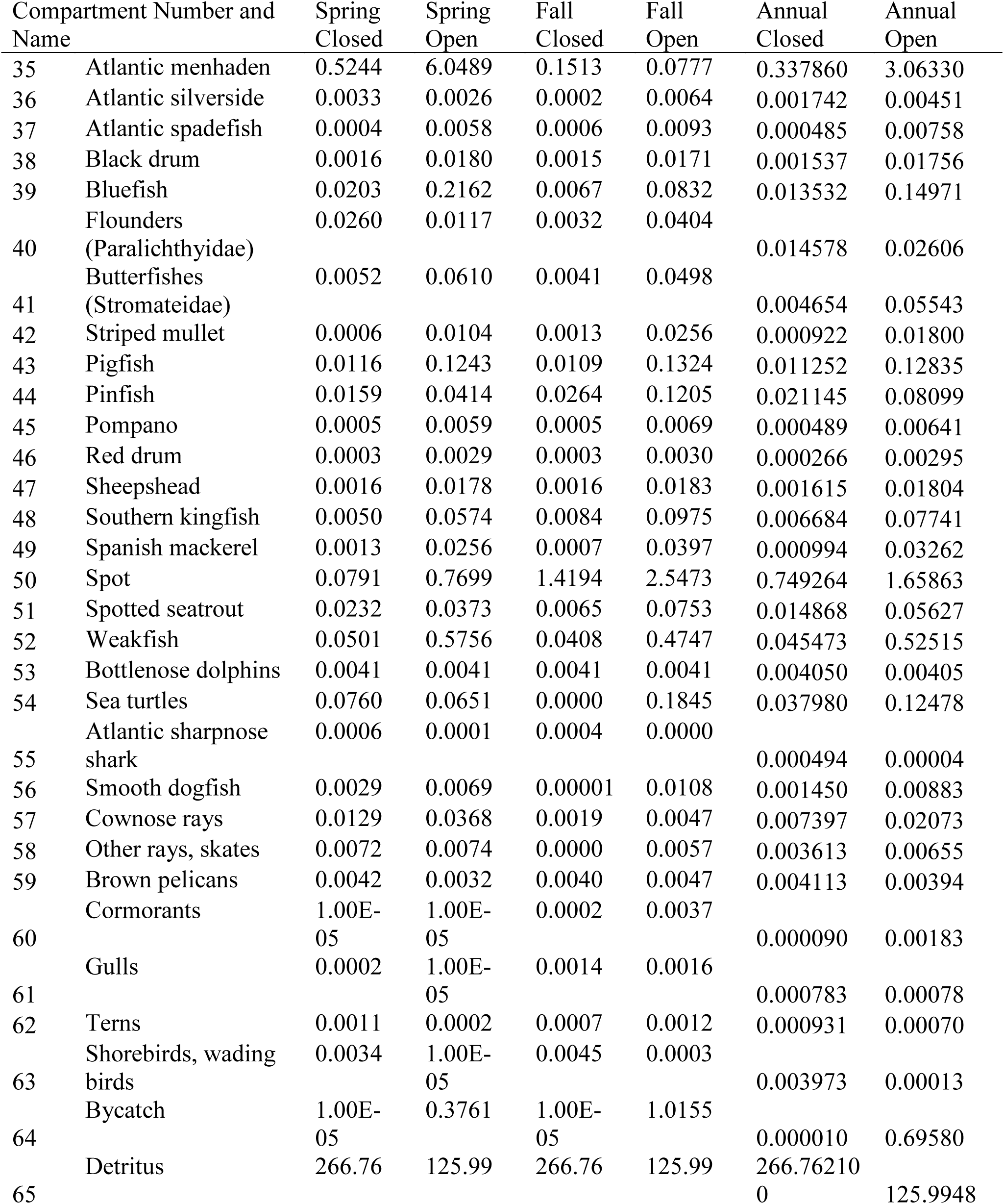
Biomass (g C m^−2^) of each compartment in the Core Sound Ecopath models. Table arranged by compartment number.

**Table 2.** Average biomass (g C m^−2^) for total macrobenthic invertebrates and the 14 sub-groups (± standard error of the mean). Averages calculated by trawling area (Open, Closed), season (Spring, Fall) and Ecopath model (Spring Closed, Spring Open, Fall Closed, Fall Open).

| Groupings | <u>Trawling Area</u> |  | <u>Season</u> |  | <u>Model</u> |  |  |  |
| --- | --- | --- | --- | --- | --- | --- | --- | --- |
|  | Closed | Open | Spring | Fall | Spring Closed | Spring Open | Fall Closed | Fall Open |
| Total macrobenthic invertebrates | 0.786 ± 0.139 | 5.044 ± 2.519 | 2.334 ± 0.714 | 3.496 ± 2.468 | 0.964 ± 0.242 | 3.704 ± 1.349 | 0.608 ± 0.133 | 6.385 ± 4.908 |
| Deposit-feeding polychaetes | 0.221 ± 0.049 | 0.712 ± 0.152 | 0.351 ± 0.086 | 0.582 ± 0.144 | 0.209 ± 0.080 | 0.492 ± 0.148 | 0.232 ± 0.059 | 0.932 ± 0.261 |
| Suspension-feeding polychaetes | 0.057 ± 0.017 | 0.100 ± 0.021 | 0.052 ± 0.012 | 0.105 ± 0.024 | 0.035 ± 0.014 | 0.068 ± 0.019 | 0.079 ± 0.031 | 0.131 ± 0.037 |
| Predatory polychaetes | 0.079 ± 0.015 | 0.217 ± 0.049 | 0.136 ± 0.037 | 0.159 ± 0.039 | 0.059 ± 0.017 | 0.214 ± 0.067 | 0.099 ± 0.024 | 0.220 ± 0.072 |
| Suspension-feeding bivalves | 0.220 ± 0.066 | 0.493 ± 0.138 | 0.562 ± 0.139 | 0.150 ± 0.050 | 0.289 ± 0.087 | 0.835 ± 0.248 | 0.150 ± 0.088 | 0.151 ± 0.051 |
| Deposit-feeding gastropods | 0.016 ± 0.006 | 0.020 ± 0.005 | 0.018 ± 0.005 | 0.019 ± 0.006 | 0.016 ± 0.007 | 0.019 ± 0.006 | 0.016 ± 0.009 | 0.021 ± 0.007 |
| Predatory gastropods | 0.047 ± 0.020 | 0.208 ± 0.150 | 0.242 ± 0.149 | 0.013 ± 0.013 | 0.093 ± 0.038 | 0.009 ± 0.003 | 0.001 ± 0.001 | 0.391 ± 0.296 |
| Amphipods, isopods and cumaceans | 0.005 ± 0.002 | 0.010 ± 0.003 | 0.007 ± 0.002 | 0.009 ± 0.003 | 0.007 ± 0.004 | 0.006 ± 0.003 | 0.003 ± 0.002 | 0.015 ± 0.006 |
| Omnivorous shrimps | 0.001 ± 0.001 | 0.001 ± 0.000 | 0.001 ± 0.001 | 0.001 ± 0.000 | 0.001 ± 0.001 | <0.000 | <0.000 | 0.001 ± 0.001 |
| Tunicates | 0.002 ± 0.002 | 2.424 ± 2.423 | 0.004 ± 0.003 | 2.423 ± 2.423 | 0.004 ± 0.004 | 0.003 ± 0.003 | <0.000 | 4.845 ± 4.845 |
| Sea cucumbers | 0.013 ± 0.013 | 0.429 ± 0.428 | 0.428 ± 0.428 | 0.014 ± 0.003 | <0.000 | 0.857 ± 0.855 | 0.027 ± 0.027 | 0.001 ± 0.001 |
| Brittlestars | 0.054 ± 0.054 | 0.338 ± 0.173 | 0.386 ± 0.179 | 0.006 ± 0.006 | 0.093 ± 0.038 | 0.009 ± 0.003 | 0.001 ± 0.001 | 0.391 ± 0.296 |
| Omnivorous crabs | <0.000 | 0.083 ± 0.059 | 0.067 ± 0.058 | 0.016 ± 0.016 | <0.000 | 0.135 ± 0.115 | <0.000 | 0.031 ± 0.031 |
| Jellyfish | 0.004 ± 0.004 | 0.011 ± 0.008 | 0.015 ± 0.009 | <0.000 | 0.008 ± 0.008 | 0.022 ± 0.016 | <0.000 | <0.000 |
| Bryozoans | 0.067 ± 0.067 | <0.000 | 0.067 ± 0.067 | <0.000 | 0.133 ± 0.133 | <0.000 | <0.000 | <0.000 |

**Table 3.**
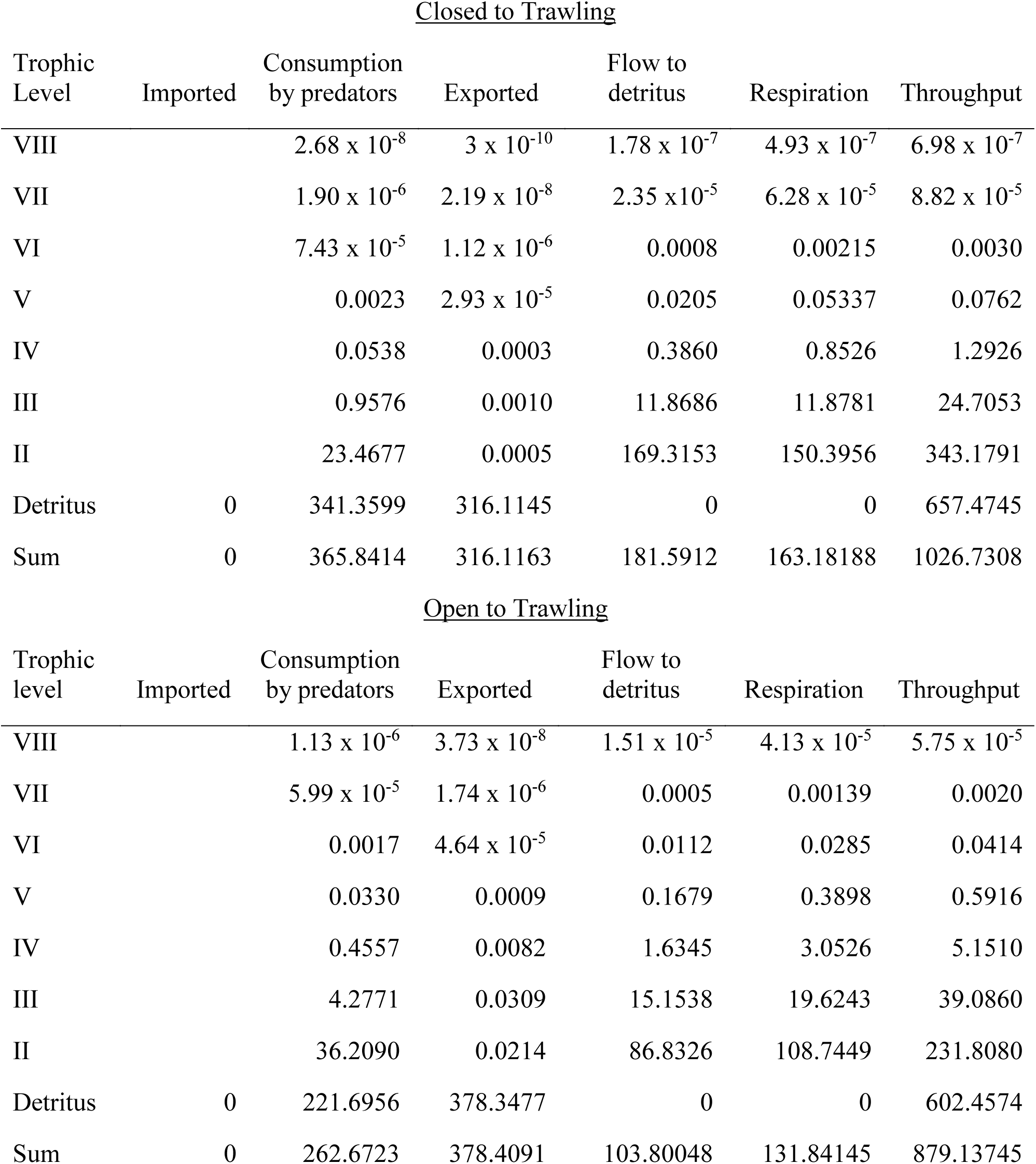
Flow of material in gC m^−2^ y^−1^ from detritus (assumed to be trophic level 1) to higher trophic levels (integrated at each level across species by Ecopath network modelling software) for the closed and open trawling areas of Core Sound, NC, USA.

To construct the Ecopath models of Core Sound, compartments encompassing everything from detritus to birds were sampled. The “currency” for these models was grams of carbon per square meter (g C m^-2^) for biomasses and grams of carbon per square meter per year (g C m^-2^ yr^-1^) for flows. For this study, biomass was measured directly for most compartments, and a diet matrix was partially constructed from the diet data obtained in Core Sound during the study period by Hart (23) for a limited number of fish species. Samples were collected in the spring, prior to the peak of commercial shrimp trawling, and then again in the fall, after the peak trawling activity ended, in areas open and closed to commercial shrimp trawling. The end result was four models, representing Spring Open, Spring Closed, Fall Open and Fall Closed. Details for all Ecopath modeling, measurements and references provided in the tables listed here are given in Deehr (24)

### Measurements of Biomass in Open and Closed Trawling Areas

Organisms’ biomasses or densities were measured at locations in the open and closed trawling areas with similar temperature, salinity, dissolved oxygen, water depth and substrate characteristics at 12 sites in Core Sound, NC (25,24) (Figure 1). We measured dry biomass converted to g C m^−2^ for all benthic groups (macrofauna and meiofauna), zooplankton, seagrasses, algae, small fishes from gill nets (three replicated nets with five 23-m panels of stretch monofilament mesh [8.9 cm, 10.2 cm, 11.5 cm, 12.7 cm and 13.9 cm] were deployed for upwards of six hours and checked at least every two hours) and bottom trawls (head rope of 3.2- m, a body net stretch mesh of 1 cm, a cod-end stretch mesh of 0.5 cm, a tickler chain, and trawl doors measuring 90 cm by 46 cm) deployed for 2 min at a constant speed, three times at each site. All biomass measurements were converted to dry weight and g C by multiplying by 0.15 (26).

**Figure 1.**
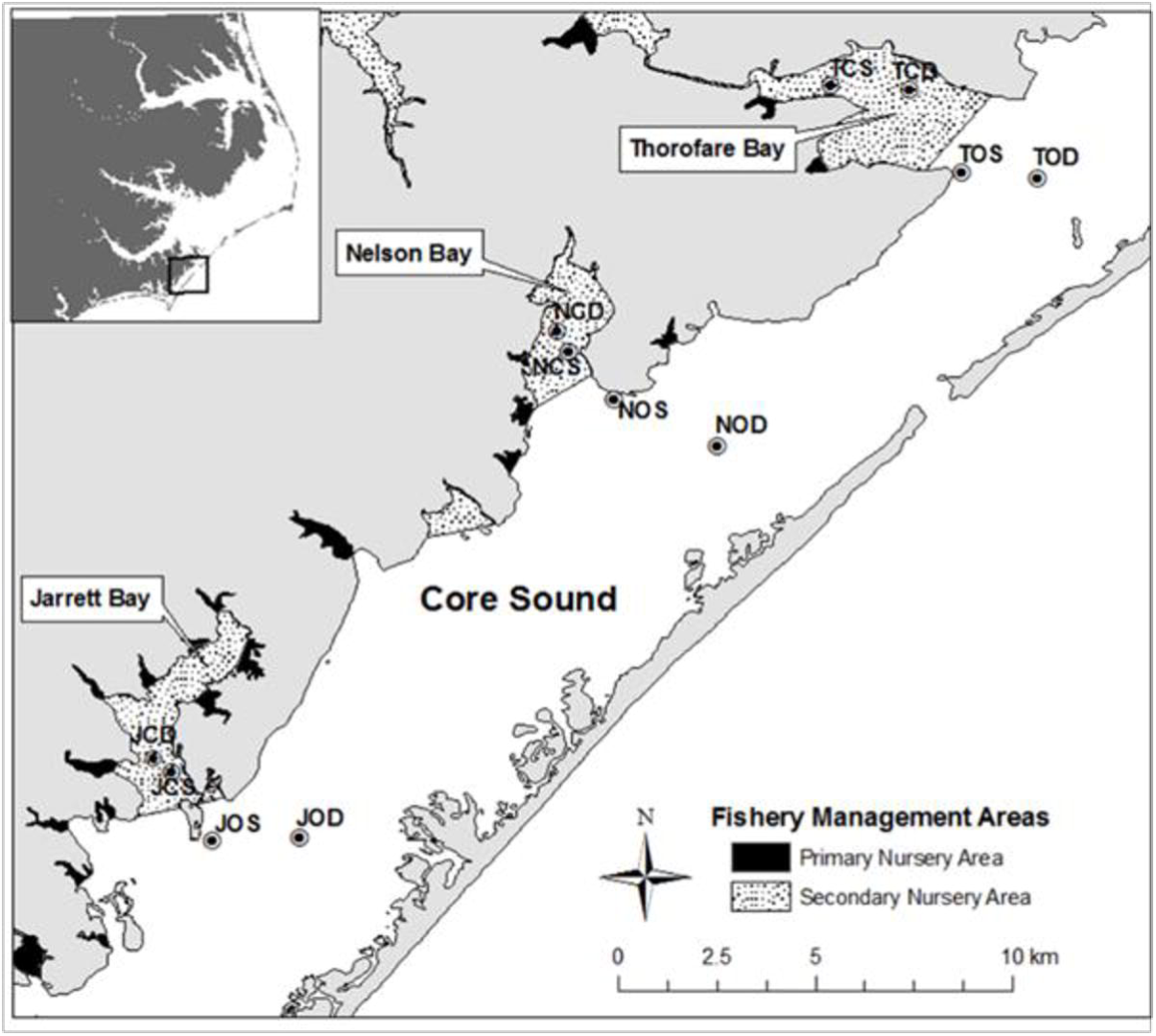
Fishery management areas in Core Sound, North Carolina, USA. Open and closed trawling areas are defined by rule (15A NCAC 03 in North Carolina Marine Fishery Commission Rules) and have been enforced for 30 years (NC Division of Marine Fisheries NCDMF). No trawling is allowed in primary and secondary nursery areas.

#### Benthic Macrofauna

At each of the 12 sites, benthic cores (inside diameter of 9.5 cm) were collected by SCUBA divers and pushed manually into the substrate to a depth of 10 cm. Three cores each were combined to form one sample that was processed for benthic macrofauna; triplicate samples were collected in this manner (a total of nine cores at each site). Three additional cores were collected at each site to obtain biomass measurements for meiofauna. Three cores each were combined to form one sample that was processed for benthic macrofauna; triplicate samples were collected in this manner (a total of nine cores). Samples were passed through a 500-μm sieve in the field, and all retained specimens were preserved in 10% buffered formalin with Rose-Bengal stain until processed in the laboratory. All specimens were identified to the lowest taxonomic level using a dissecting microscope, then dried at 60°C for 48 hr, weighed to the nearest 0.00001 g, then converted to carbon by multiplying dry weight by 0.40(26).

#### Benthic Meiofauna

The remaining three benthic cores were sub-sampled for meiofauna, detritus, benthic microalgae and sediment grain size. Meiofauna were collected from each core with a 2-cm diameter syringe plunged to a depth of 3 cm, and preserved in 10% buffered formalin with Rose-Bengal stain. Meiofauna were separated from sediments using Ludox, following the method of Burgess (2001), passed through stacked 500-μm and 63-μm sieves (to exclude macroinvertebrates), and all specimens retained on the 63-μm sieve were identified to lowest taxonomic level using Higgins and Thiel (27). All individuals (by taxa) were converted to g C from wet weight/individual and/or g C/individual from several sources (24,28).

#### Benthic Detritus

Detritus samples were collected from each core using a 1-cm diameter syringe pushed to a depth of 1 cm and stored on ice in a dark cooler then frozen until processed in the laboratory. Loss on ignition (LOI) was used to determine the ash-free dry mass of organic matter(29). Since the sample potentially included numerous sources of organic matter, values of sediment microalgae, sediment bacteria and meiofauna biomasses (also calculated for this study) were subtracted from the LOI-obtained measurement of organic carbon. Dry weights were converted to g C by multiplying by 0.58(26).

#### Benthic Microalgae

Similarly, benthic microalgae biomass was sampled from each core using a 1-cm diameter syringe plunged to a depth of 1 cm. Benthic microalgal biomass was measured using fluorometry as the amount of chlorophyll *a* content in the sample. Chlorophyll *a* was converted to g C by multiplying by 0.47(26). Only the samples collected during the spring (for open and closed sites) were processed in the laboratory; thus, there is no seasonal difference between benthic microalgae biomass.

#### Infaunal Mollusks

A clam rake was used to collect mollusks from sites in shallow water. Four 2.32-m^2^ transects (total area of 9.29 m^2^) were raked at the six shallow sites. Mollusks were stored on ice until returned to the laboratory for positive identification and measurements. All specimens were removed from the shells and dried in an oven at 60°C for 48 hr. Dry weight mass was converted to g C by multiplying by 0.40(26).

#### Benthic Primary Producers

The biomass of primary producers (macroalgae, drift algae, seagrasses) was measured using various techniques. Seagrass biomass estimates were obtained from an ongoing submerged aquatic vegetation study in Jarrett Bay using 0.15-m cores and quadrats, as well as remote sensing, and video and acoustic methods. Data from a site in the area closed to trawling in Jarrett Bay were collected from June – September 2010; thus, seagrass measurements for this project differ by season, but not by area. The values for seagrass biomass are only from the closed areas, but also used for the open areas of the Core Sound models. Drift algae and macroalgae biomass estimates were calculated from algae collected in otter trawls for sites open and closed to trawling, but data were only collected in the fall. Thus, there are no seasonal differences in biomass for the models (the same values for fall were used for spring). Otter trawl distances were obtained from a digital echo-sounder (see Nekton below).

#### Zooplankton

Three replicates of zooplankton samples were collected at each site using 90-μm mesh bongo plankton nets (net diameter of 28 cm), towed for 1 min at a constant speed. Continuous GPS locations throughout the tows were recorded to avoid crossing previous tow tracks and to obtain the tow distances. A General Oceanics flow meter with the low-speed rotor was attached to the bongo net to measure the volume of water towed. All zooplankton samples were fixed in 10% buffered formalin for storage until processing. Any ctenophores or other large gelatinous zooplankton were removed before fixing. To estimate the abundance of ctenophores, separate 1- min tows were conducted. Any ctenophores collected in the tows were counted and recorded on the boat. Total counts of ctenophores were converted to biomass [assumed one ctenophore had wet weight of 1 g, multiply by 0.20 to convert to dry weight then g C] for use in the Ecopath model. In the laboratory, all large zooplankton specimens (≥ 500 μm) were counted and dried at 60°C for 48 hr to measure dry mass, which was converted to g C by multiplying by 0.40. Using a Folsom splitter, the remainder of the samples were split three times, and the 1/8 sample was suspended in 500 mL of water. Five 10-ml subsamples were taken with Hensen-Stempel pipettes and passed through a series of sieves (425 μm, 250 μm, 150 μm, and 75 μm). The contents of each sieve were counted in a Ward wheel, identified to lowest taxonomic level, summed and total counts were multiplied by 80 to obtain the whole sample count. This method subsampled at least 100-300 individuals at a time, an amount recommended by several sources to avoid potential errors associated with repetitive Folsom splitting of samples. The entire contents of each sieve were dried at 60°C for 48 hr to calculate dry mass, and then converted to g C by multiplying by 0.40 (26).

#### Phytoplankton

Water samples were collected at each site to measure phytoplankton. Carboys (1 L^3^) were filled with surface water at each station and stored on ice in a cooler until returned to the laboratory. In the laboratory, water was filtered through glass microfiber filters (47 mm, GF/C). Pigment extraction was done with a mixture of 45% acetone/45% methanol/10% deionized water, then kept in a freezer for 12-24 hr, using the methods of Strickland and Parsons (30). Initial readings were done on the fluorometer, then 10% HCl was added, to correct for pheophytin pigments, and then read again. Chlorophyll *a* values were then converted to g C by multiplying by 0.47(26).

#### Nekton

To sample fishes and other forms of nekton, an otter trawl similar to the one used by NC DMF was deployed. The protocols that follow are from Hart (23). The otter trawl had a headrope of 3.2-m, a body net stretch mesh of 1-cm, a cod-end stretch mesh of 0.5-cm, a tickler chain, and trawl doors measuring 90 cm by 46 cm. Trawls were deployed for 2 min at a constant speed, three times at each site. Trawl tow lengths were determined using a scientific echo-sounder operated simultaneously with the trawl deployment. The BioSonics DTX echo-sounder was used to assess bathymetry, bottom substrate, and fish abundance in front of the trawl. The echo-sounder was interfaced with a JVC GPS receiver and a Panasonic Toughbook CF-29 laptop computer so that precise trawl tracks and depths were recorded to a hard drive (23). All specimens retained by the trawls were euthanized and preserved in 10% buffered formalin for identification and measurement in the laboratory. When necessary, some samples were weighed in the field using spring scales. In the laboratory, all specimens were identified, measured for length and wet weight, and stomachs of selected fishes were removed for diet analyses. All biomass measurements were converted to dry weight and g C by multiplying by 0.15 (26).

Experimental gill nets were used to collect larger, faster fishes not captured by the otter trawl. Five 23-m panels of different stretch mesh (8.9 cm, 10.2 cm, 11.5 cm, 12.7 cm and 13.9 cm) were deployed for upwards of six hours and checked at least every two hours. All specimens were euthanized, tagged and stored on ice in a cooler until brought back to the laboratory or field processing site. Specimens were identified, measured and stomachs were removed for diet analyses. All biomass measurements were converted to g C by multiplying by 0.15 (26).

Additional fish and shellfish data were obtained from the North Carolina Division of Marine Fisheries (NCDMF) Program 120 Juvenile Trawl Survey (Katy West, personal communication, NCDMF, 3441 Arendell St, Morehead City, NC 28557 USA). Trawl surveys have been conducted in the spring in nursery areas to inform management decisions on the opening and closing dates of various fisheries. Data for several species of fish and shrimps were included in the construction of the models in this study.

#### Fisheries Data

Unpublished NCDMF Trip Ticket data from April-June 2006 and 2007 (averaged to represent Spring) and August-October 2006 and 2007 (averaged to represent Fall) for the six fishing gears described in Chapter One (shrimp trawls, skimmer trawls, pound nets, crab pots, haul seines and gill nets) were included in the models for this study. If a landings report was made but unavailable (due to confidentiality), an average of years 2001-2005 for that gear type, species and month (in 2006 or 2007) was used. The area of Core Sound waters was estimated to be 72,000 acres (291,272,662 m^2^). The average catch (in wet weight pounds) was converted to grams of wet weight then multiplied by 0.15 (26) to convert g C dry weight, and finally divided by the area of Core Sound (resulting in g C/m^2^ for each species by gear type). Because trawlers cannot operate in closed areas or in known seagrass beds, the area for calculating shrimp trawl and skimmer trawl catches was reduced by 50% (145,686,831 m^2^). These values represent the biomass of each species that was added to our own data collections (from juvenile trawls and gill nets). To calculate fisheries trip averages (for fisheries landings data in Ecopath), we used the pounds/trip average of the time periods listed above and calculated g C for those data. To convert fisheries trip averages to g C/m^2^, we estimated the area fished by each gear type, based on our knowledge of the gears, the information provided by the NCDMF, and shrimp trawl and skimmer trawl bycatch studies (**Error! Reference source not found.**).

Information about shrimp and skimmer trawl landings were incorporated only in the models representing areas open to trawling. Data about the landings of the other four gears were split 10% in the Closed models and 90% in the Open models, based on the relative areas of closed and open waters in the study, respectively.

Bycatch from trawls was also included in the models for this study. Bycatch data for shrimp and skimmer trawls were available from local studies conducted in and near Core Sound (17,22,31). These data are included in (**Error! Reference source not found.**). While bycatch is known to occur with the other four gear types, studies reporting bycatch statistics for gill nets, pound nets, haul seines and crab pots were insufficient for inclusion in this study.

Trip ticket landings data were organized using six fishery gear types (crab pots, haul seines, pound nets, gill nets, skimmer trawls and shrimp trawls) that were included in the Ecopath models as fishing fleets. Inclusion of the NCDMF Trip-Ticket data in the Ecopath model required us to increase the biomass in the open trawling areas relative to the closed trawling areas, because most of the trip-ticket landings in Core Sound were reported from trawling gear. We determined that >90 % of the commercial harvest in Core Sound came from open trawling areas. Because trawling gear is not allowed in closed trawling areas (NCDMF designated Primary or Secondary Nursery Areas) all trip-ticket data from trawling gear was included in the open-trawling Ecopath model only. In contrast, for other fishing gear types (pound nets, gill nets, haul seines, and crab pots), which can be used in either open-trawling areas and the Secondary Nursery Areas (but not Primary Nursery Areas), we made the assumption that 90% of the Trip-Ticket catch (in pounds converted to Ecopath biomass per unit area g C m^−2^) originated in the open trawling areas, based on the relative size of Secondary Nursery Areas (10% of the total area of Core Sound). Biomass estimates from some of our field measurements were insufficient to account for reported landings, as indicated by Ecotrophic Efficiencies > 1 in Ecopath, and the carbon energy required by the commercial fisheries of Core Sound could not be met by production of lower trophic levels.

### Ecosim simulation modelling

Ecosim is a module of Ecopath that allows time-dynamic simulations of balanced Ecopath models (38). EcoSim was run after balanced EcoPath models were achieved. Default settings for vulnerability were used (2% of each prey population was available for predator consumption at any given time), and the system was calibrated with historical catch and effort data. The Ecopath model used was the Annualized Open trawling model for the simulation (SI Tables 4-8). We drove the model effort statistics for each year for the fishing fleets (Table SI 10) as reported on Trip Tickets to the NCDMF for the shrimp trawling fishery in the Core Sound Management Area for the years 2001-2007. The model was fit to time series of annual catches of brown shrimp, white shrimp, pink shrimp, blue crabs, flounders, spot, pinfish, and other species for each fish gear (Table SI 9). In Ecosim, vulnerabilities are parameters that assign a value for each species or node indicating the proportion of that node’s population that is available to preyed upon by other consumers. A vulnerability of 0.0 would indicate that all biomass in that node would be immune from any predation, whereas V = 100.0 or greater would indicate that all the individuals and their biomass are vulnerable to predation. The vulnerabilities were kept initially at the default values of 2.0. The vulnerabilities are essential in maintaining the model within stable boundaries, and provide a degree of refugia for each of the nodes. We used the Ecosim module “fit to time series” to estimate vulnerability parameters that minimize the sum of squares between predicted and observed values (SS= 112.1313, Akaike Information Criterion, AIC= 435.5736). Our model effort time series reflected the real trawling and other fishing gear relative fishing effort during the historical time period for which harvest data were available, and this served as a basis for simulating the trawling effort reduction. Next, a “trawl ban” was simulated by setting the shrimp trawling effort = 0 trips per month for the years 2008-2026. Other gears were left with a relative fishing effort of 1.0 over this same period, (i.e., effort levels as reported in 2007, the base year for which our Ecopath models were developed). The runs of the model with the “trawl ban” were reported for 18 years after trawling effort ceased and are reported here.

**Table 4.** The species identification codes for food web visualizations shown in Figure 5 and Figure 6.

| Code on graph | Ecopath ID | Group name in Ecopath |
| --- | --- | --- |
| 1 | 26 | Amphipods, isopods, cumaceans |
| 2 | 33 | Anchovies |
| 3 | 21 | Atlantic brief squid |
| 4 | 34 | Atlantic croaker |
| 5 | 35 | Atlantic menhaden |
| 6 | 55 | Atlantic Sharpnose shark |
| 7 | 36 | Atlantic silverside |
| 8 | 37 | Atlantic spadefish |
| 9 | 6 | Bacteria aquatic |
| 10 | 7 | Bacteria benthic |
| 11 | 16 | Bay scallop |
| 12 | 15 | Bivalves, suspension feeding |
| 13 | 38 | Black drum |
| 14 | 27 | Blue crabs |
| 15 | 39 | Bluefish |
| 16 | 53 | Bottlenose dolphins |
| 17 | 25 | Brittlestars |
| 18 | 59 | Brown pelicans |
| 19 | 29 | Brown shrimp |
| 20 | 22 | Bryozoans |
| 21 | 64 | Bycatch |
| 22 | 20 | Conchs, whelks |
| 23 | 60 | Cormorants |
| 24 | 57 | Cownose rays |
| 25 |  | Crab Pots |
| 26 | 28 | Crabs other |
| 27 | 11 | Ctenophores |
| 28 | 65 | Detritus |
| 29 | 4 | Drift algae |
| 30 | 40 | Flounders (Paralichthyidae) |
| 31 | 18 | Gastropods, deposit feeders |
| 32 | 19 | Gastropods, predatory |
| 33 |  | Gill Nets |
| 34 | 61 | Gulls |
| 35 | 17 | Hard clams |
| 36 | 41 | Butterfishes (Stromatiedae) |
| 37 |  | Haul Seines |
| 38 | 10 | Jellyfish |
| 39 | 3 | Macroalgae, benthic |
| 40 | 8 | Meiofauna |
| 41 | 2 | Microalgae, benthic |
| 42 | 58 | Other rays and skates |
| 43 | 1 | Phytoplankton |
| 44 | 43 | Pigfish |
| 45 | 44 | Pinfish |
| 46 | 30 | Pink shrimp |
| 47 | 12 | Polychaetes, deposit feeders |
| 48 | 14 | Polychaetes, predatory |
| 49 | 13 | Polychaetes, suspension feeders |
| 50 | 45 | Pompano |
| 51 | 46 | Pound Nets |
| 52 | 24 | Red drum |
| 53 | 10 | Sea cucumbers |
| 54 | 54 | Sea turtles |
| 55 | 5 | Seagrass |
| 56 | 47 | Sheepshead |
| 57 | 63 | Shorebirds/waders |
| 58 |  | Shrimp Trawls |
| 59 | 32 | Shrimps other |
| 60 |  | Skimmer Trawls |
| 61 | 56 | Smooth dogfish |
| 62 | 48 | Southern kingfish |
| 63 | 49 | Spanish mackerel |
| 64 | 50 | Spot |
| 65 | 51 | Spotted seatrout |
| 66 | 42 | Striped mullet |
| 67 | 62 | Terns |
| 68 | 23 | Tunicates |
| 69 | 52 | Weakfish |
| 70 | 31 | White shrimp |
| 71 | 9 | Zooplankton |

### Visualization of the Trophic Networks

To visualize the food web, we plotted each of the 65 nodes plus the 6 fleets using the log-10 transformed biomass (or catch) of each node and the consumption (*Q_i,j_*) matrices from Ecopath, where prey *i* is consumed by predator *j*, and the consumption in gC m^−2^ yr^-1^ of *i* by *j* is given in each cell. Consumption matrices for each model from Fall 2007, areas open and closed to trawling, were analyzed for similarity in trophic roles using regular equivalence (REGE, (32,33) as a measure of similarity. The REGE coefficients were plotted in a 2-D multidimensional scaling coordinates with in UCI Net 6.361 and Pajek64 3.12 (34,32,35). The algorithm takes any real-valued *N* × *N* (species-by-species) matrix *X* as input, and returns a species-by-species matrix *R* of coefficients (ranging from 0 to 1) which records, for each pair of species, the extent of (maximal) regular equivalence. The essence of the algorithm is as follows:

0. Set r_ij_ = 1 for all i and j (i.e., let all species be 100% equivalent to start)
1. For each species i and j,

A. For each species *k* eaten by *i*, find species *m* eaten by *j* that is most equivalent to *k* and which is eaten in the most similar proportion as *k* is eaten by *i*, in other words, which maximizes the quantity *z*_*k*_ = r_km_^*^Min(x_ik_,x_jm_)/Max(x_ik_,x_jm_)
B. For each *k* which eats *i*, find species *m* that eats *j* that is most equivalent to *k* and which eats *j* in the most similar proportion as *k* eats *i*, in other words which maximizes the quantity *y*_*k*_ = r_km_^*^Min(x_ki_,x_mj_)/Max(x_kj_,x_mj_)
C. Set r_ij_ and r_ii_ = Σ *z*_*k*_ + Σ *y_k_*
2. Repeat Step 1 until no more changes in r_ij_ or maximum iterations exceeded. The maximum iterations = *N* species or compartments.

The resulting coefficients *r*_*ij*_ have ordinal properties.

Trophic roles are thought to be most similar in this analysis when the REGE coefficient r_ij_ is large between any pair of nodes, indicating a similar trophic niche (predator’s with similar trophic roles or niches and prey with similar trophic roles or niches, but not the exact same predator or prey). The REGE algorithm is iterative; a minimum of 50 iterations were used to obtain the REGE coefficients for each trophic network model, and as nodes were assessed for trophic similarity at each iteration, the REGE coefficients from the previous iteration were used to assess trophic similarity in the next iteration. All nodes begin in one group at the first iteration (REGE is set at 100 % similarity for all nodes), and trophic role similarity was used to establish the REGE coefficients at each iteration of the algorithm, finding the nodes that are least similar to the group and giving them a new, lower REGE coefficient.

After running the REGE algorithm on each of the seasonal and trawling area closure networks (65 Ecopath compartments plus 4 closed trawling area or 6 open trawling area fishing gears as nodes in the network), a clustering strategy was applied [Johnson’s hierarchical clustering strategy in UCINet (36)] to the resulting matrix (71 × 71 node by node matrix for open trawling areas and 69 × 69 nodes for closed trawling areas) of REGE coefficients. Because it is appropriate for ordinal data, we used Johnson’s hierarchical linkage clustering, which yields a dendrogram and a set of nested partitions.

To simplify interpretation of the results, a hierarchical clustering of the output matrix E from the REGE algorithm was also performed, yielding a dendrogram. For display purposes, one partition within the hierarchical clustering was selected to classify compartments. The particular choice of partition was based on a series of regressions designed to measure cluster adequacy. Since an ideal clustering of the E matrix would locate the largest values of E within clusters and the smallest values of E between clusters, we can measure the extent to which a given clustering is optimal via an analysis of variance in which the cases are pairs of nodes, the dependent variable is the REGE coefficient for each pair, and the independent variable is a dummy variable coded 1 if the pair are in the same cluster and 0 if they are in different clusters. The resulting R-square (or η^2^ as it is called in the ANOVA context) is then interpreted as a measure of cluster adequacy. By necessity, R-square is a non-decreasing function of the number of clusters. By plotting R-square against the number of clusters we obtain a scree plot which can be examined for inflection points. A clustering with k classes is chosen if it provides a sizeable increase in R-square over the next simplest clustering (i.e., with k-1 clusters), yet explains nearly as much variance as the next most complicated clustering (k+1 clusters).

The cluster adequacy scree plots [η^2^ plotted versus cluster partition group size; η^2^ is a measure of within-cluster versus between-cluster variance (37)] was plotted for the four models are shown in Figure 2. The plot shows that when all nodes are grouped as one large cluster (on left side of plot), η^2^ is low. Conversely, when each node is assigned to its own individual cluster partition group (resulting in 69 - 71 clusters, on the right side of the plot), η^2^ is also very low. When η^2^ is maximal, the clustering partitioning is most adequate at capturing the within group variance in REGE coefficients. The four models had slightly different η^2^ maxima for number of cluster groups (Spring Open η^2^= 0.75, Spring Closed η^2^=0.75, and Fall Closed η^2^ = 0.76 occurred at 15 clusters; Fall Open was maximal η^2^ = 0.75 at 9 clusters), and these groups with maximal within group REGE coefficients were used to assign color classes to nodes with high REGE similarity.

**Figure 2.**
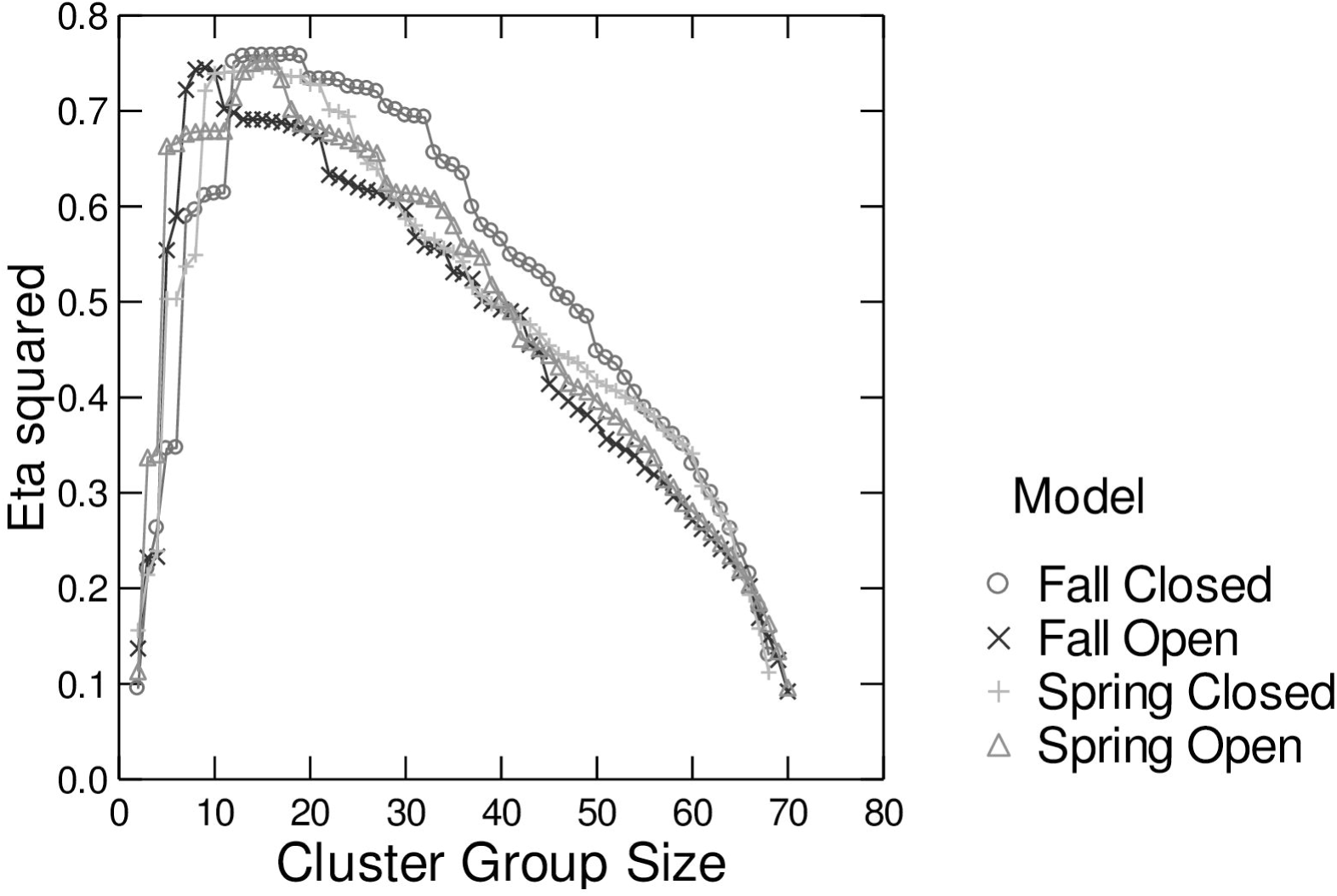
Core Sound networks clustering results. Scree plot of cluster group size and η^2^ for each of the Core Sound models, based on Johnson’s hierarchical clustering procedure of the REGE coefficients from UCINET of the four Core Sound network models.

In food web these visualizations, if two nodes have similar REGE coefficients, they are likely to have similar trophic roles and will plot near one another on the MDS coordinates. Node size on each food web visualization was scaled by log-10 biomass: in addition, we plotted the difference in log-10 biomass measured between open and closed areas in fall (2007), thus providing node-by-node a ratio of the open: closed biomass, and closed: open biomass.

## Results

### Benthic Biomass in the Closed and Open Trawling Areas

In Core Sound, North Carolina, shrimp trawling starts in March and runs through October (39). After the peak of the shrimp trawling season in the fall of 2007, benthic deposit-feeding polychaetes biomass (Figure 3) was higher in the open trawling areas than in closed trawling areas [Table 1, repeated-measures ANOVA between trawling areas: total macrobenthic invertebrates (F_1,34_ = 6.210, p = 0.018), deposit-feeding polychaetes (F_1,34_ = 7.894, p = 0.008) and predatory polychaetes (F_1,34_ = 6.339, p = 0.017)]. Deposit-feeding polychaetes are scavengers and consume dead fishes, organic material and smaller bacteria and microbes.

**Figure 3.**
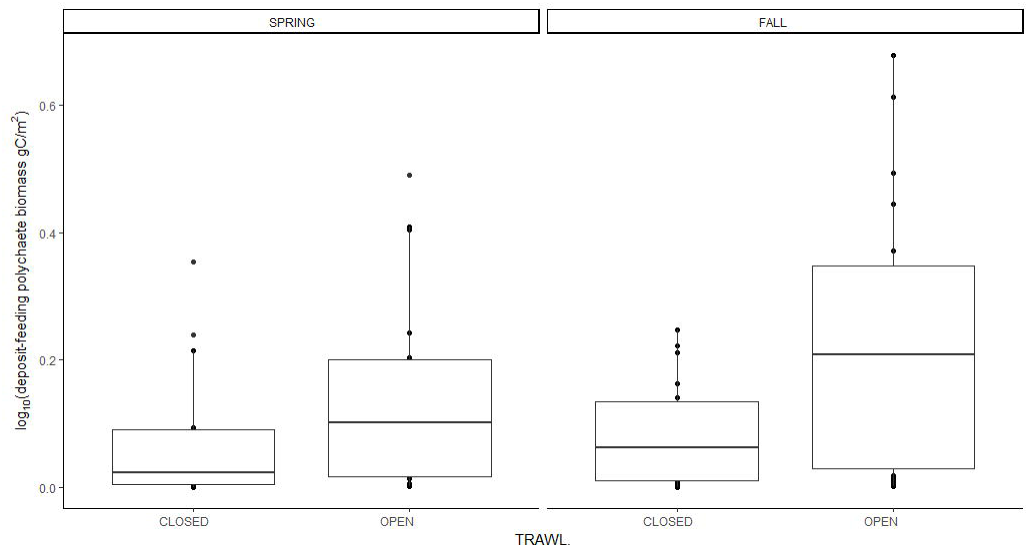
The log-transformed biomass m^−2^ of deposit-feeding polychaetes in Core Sound, NC in areas closed (nursery areas) and open to shrimp trawling. Individual points show the abundance of polychaetes at each of stations sampled during 2007, before (Spring 2007) and after (Fall 2007) the shrimping season.

### Bycatch Biomass in the Closed and Open Trawling Areas

The three species most commonly caught as bycatch in shrimp trawls are blue crabs (*Callinectes sapidus*, Portunidae), pinfish (*Lagodon rhomboides*, Sparidae), and spot (*Leiostomus xanthurus*, Sciaenidae) (17). These were collected in our own replicated trawl samples (using a smaller version of the otter trawls used by shrimpers, with smaller mesh liner and shorter headrope) to capture these bycatch species at the start of and after the shrimp season (spring and fall 2007) at the same stations as the benthic samples were taken above. There was significantly greater biomass of the three main bycatch species in the closed trawling areas at the end of the trawling season (Figure 4, Wilcoxon text, blue crab: W = 945, p>0.00001, pinfish: W=832, p> 0.04, spot: W=1062, P> 0.00001, n=36 trawls/trawling area in each case). Stomach content analysis and stable isotope estimates of the spot and pinfish diets showed that they consumed predominantly polychaetes, among many other invertebrate prey, algae and plants. Thus, predation by these bycatch fish species on deposit-feeding polychaetes was likely to be far lower in the open trawling areas, especially at the end of the shrimp trawling season, and these data were included in the Ecopath models that we constructed next.

**Figure 4.**
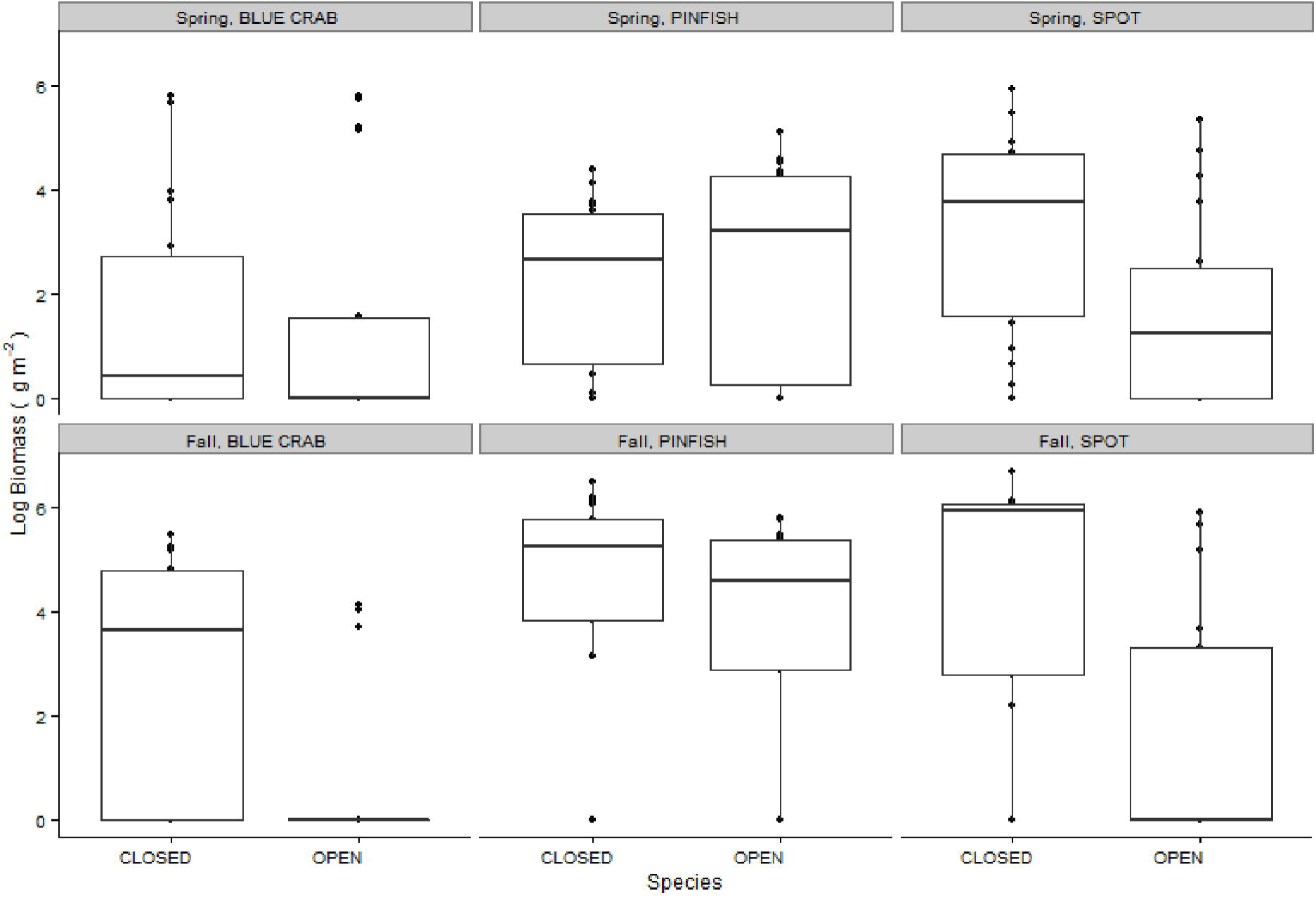
The main bycatch species (blue crabs, *Callinectes sapidus*, pinfish, *Lagodon rhomboides*, and spot, *Leiostomus xanthurus*) log_10_ biomass (g m^−2^) estimates in open and closed trawling areas before shrimp season has begun (Spring) and after it is over (Fall). The box plots show the median (horizontal lines), 25% and 75% percentiles (upper and lower limits of the box), and the whiskers are 1.5 ^*^ the inter-quartile (distance between the upper and lower box limits). Points outside the whiskers are extreme values.

### Ecopath and Food Web Model of Core Sound

The food web network models of the open and closed shrimp trawling areas of Core Sound after the shrimping season was over (in the fall months) showed a dramatic change in the benthos (Figure 5). The trophic levels of all compartments in open and closed areas are given in supplementary information (species names Table SI-2; dietary sources in SI-3; effective trophic levels Table SI-4; biomass values in Table SI-5; production/biomass ratios Table SI-6; consumption/biomass ratios in Table SI-7; ecotrophic efficiencies in Table SI-8). To make these visualizations of the Ecopath network models, we used a method based on graph theory (regular equivalence algorithm, or REGE (37)) to assess the trophic role similarity in each fishery management area. These food web flow diagrams are based on a two-dimensional multidimensional scaling of the nodes, and thus nodes with similar REGE coefficients (and trophic roles) plot close to one another, with high similarity indicated by the same color class (no two species had identical REGE coefficients; color classes with high-within class REGE similarity were determined using a clustering algorithm (Figure 2), along with species ID codes (Table 4). The position of each node in the vertical and horizontal dimensions of these plots is interpreted as depicting a trophic role for each species that is influenced not only by the relationship to the producers (39, 43, 55) and detritus (28) at the bottom of the plots, but also their relationships to their predators. Thus apex predators, including the various fishery gears [crab pots (25), gill nets (33), haul seines (37), pound nets (51), shrimp trawls (58), and skimmer trawls (60) appear near the top of the plots.

**Figure 5.**
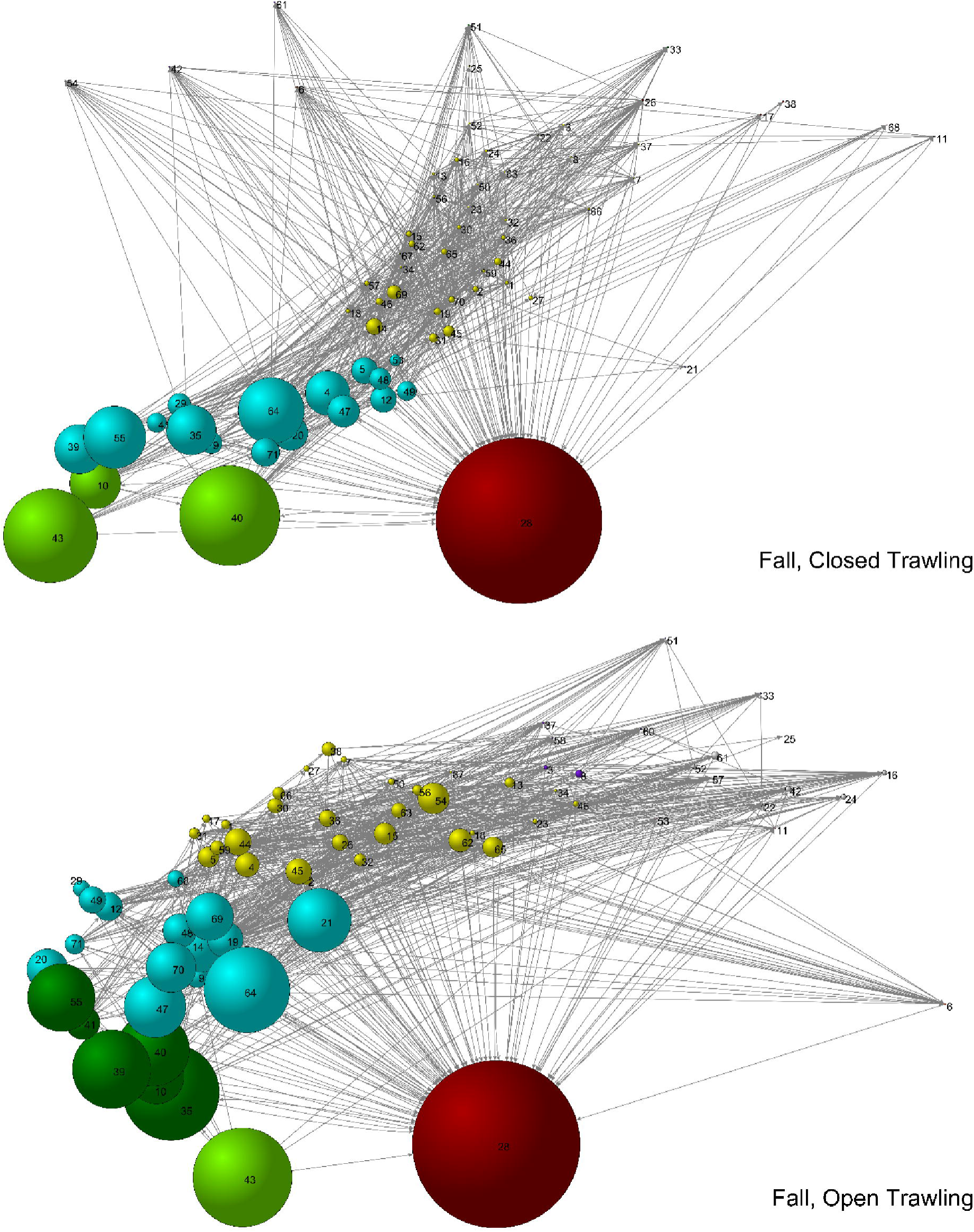
The ecological network model of Core Sound shrimping areas, with nodes representing standing stocks (size of node proportional to biomass in C dry mass g m^−2^) and flows of carbon (gC m^−2^ yr^−1^) shown as arrows. Nodes are arranged according to their similarity of trophic niche, using a non-metric multidimensional scaling (MDS) of the REGE coefficients. Nodes with the same color are similar in trophic niche (>90% similar as judged by a REGE coefficient See clustering results in SI). Top: areas closed to trawling in the fall, after shrimp season (MDS stress=0.099 after 42 iterations); Bottom: areas open to trawling after shrimp season (MDS stress=0.091 after 47 iterations).

The nodes in Figure 5 are scaled by log_10_ biomass (g C/m^−2^). The food web of Core Sound is detritus-based. Detritus (28) was considered a non-living compartment and designated trophic level 1 in the Ecopath modelling approach; this very large-biomass node appears at the bottom and in a central position on the flow diagrams in Figure 5. A general decrease in biomass is apparent as the trophic position of each species increases, with species apex predators such as various fishes [southern flounders (*Paralichthys lethostigma*) and other Paralichthidae (30), bluefish (*Pomatomus saltatrix*, 15), red drum (*Sciaenops ocellatus*, 52), sharks and rays (6, 25, 42, 61), birds (18,57,67) and sea turtles (54) having small biomasses and plotting near the top of flow diagram. Producers [seagrasses (55), phytoplankton (43), and benthic macroalgae (39) have large biomasses and plot near the bottom of the diagram. Note that biomass of bycatch (21) is small in the closed areas (from some legal fisheries in the no-trawling management areas), but a very large amount of bycatch biomass is present in the open areas. This bycatch biomass is the basis of a scavenger food web [the benthic bacteria (10), meiofauna (40), deposit-feeding polychaetes (47), and indirectly the blue crabs (14).

After the shrimp trawling season was largely over, in the fall of 2007, more detritus was found in the closed areas than in the open areas of Core Sound. Flows of C in Core Sound were dominated by consumption of detritus by benthic bacteria (10), meiofauna (40), and higher trophic levels species (Table 2). More C flowed from detritus to all predators in the closed area (365.84 gC m^−2^ yr^−1^) than in the open trawling areas (262.67 gC m^−2^ yr^−1^). Most of this flow is from detritus to consumers at Trophic level II (i.e. detritivores). Most modeled compartments had greater biomass in the open trawling area after the end of the shrimp season (Figure 6, top) including bluefish (*Pomatomus saltatrix*, 15), weakfish (*Cynoscion regalis*, 69), spotted seatrout (*C. nebulosus*, 65), Spanish mackerel (*Scomberomorus maculatus*, 63), Atlantic menhaden (*Brevoortia tyrannus*, 5), spot (64), pinfish (45), hard clams (*Mercenaria mercenaria*, 35), suspension feeding bivalves (12), blue crabs (*Callinectes sapidus*, 14), brown (*Farfantepenaus aztecus*, 19), pink shrimp (*F. duorarum*, 46), white shrimp (*Litopenaeus setiferus*, 70), polychaetes (47–49), sea cucumbers (53), and brittlestars (17). In contrast, detritus (28), drift algae (29), meiofauna (40), phytoplankton (43), zooplankton (71), and Atlantic croaker (*Micropogonias undulatus*, 4) have more biomass in the areas closed to trawling (Figure 6, bottom).

**Figure 6.**
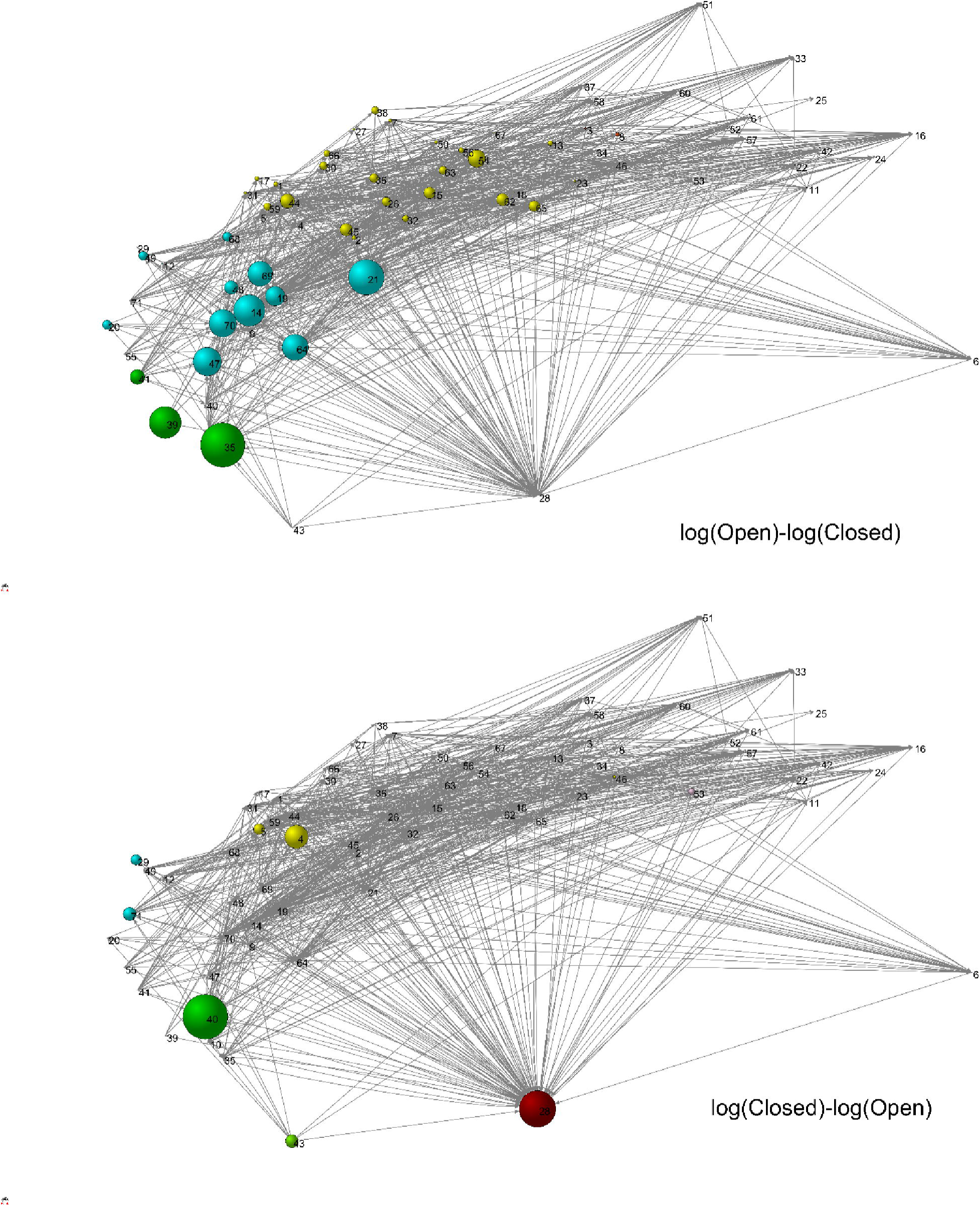
The Core Sound food web, with nodes arranged using MDS as in Figure 2 bottom (Fall, Open), but with node size scaled as the difference in log_10_ biomass between open trawling and closed trawling areas (i.e., the ratio of biomasses in the two areas). Top graph: nodes in which (log10 open biomass - log10 closed biomass) > 0, i.e., where the biomass in open trawling areas exceeded closed trawling areas; Bottom graph: nodes in which (log_10_ closed biomass – log_10_ open biomass) > 0, i.e., where the biomass in closed trawling areas exceeded open trawling areas.

The node-specific log-biomass differences (ratios) displayed in Figure 6 are based on single biomass estimates used to create balanced stead-state Ecopath models. For each species or node, a biomass was estimated from all of our samples and the North Carolina Division of Marine Fisheries (NCDMF) harvest data, one estimate/node for open trawling areas and one estimate/node for closed trawling areas. The steady-state network models, balanced to achieve steady-state conditions (see SI for method used for steady-state model balancing), were then used to perform Ecosim simulations. Biomass estimates for fishery species are based on modelled parameters derived from fisheries harvest data reported to NCDMF, and subdivided into the open and closed areas (e.g., no commercial trawl harvest data were assigned to closed areas, but other gear types were allowed in the closed trawl areas; these were proportionally divided by relative amount of fished areas), which means that there are no statistical uncertainties associated with these estimates. For biomasses of the benthos, where these were directly measured with replication, see Table 1 and results displayed Figure 3.

### Simulation modelling in EcoSim

Ecopath was used to simulate the closure of Core Sound to shrimp trawling using the EcoSim simulation module. We used an Ecopath open trawling area annual model (Deehr *et al*. 2014) that was verified with stable isotope measurements and included a time series of fisheries harvest data for Core Sound (SI Table 9) to calibrate the Ecosim model along with fishing effort (annual trips SI Table 10) by gear type reported to the NCDMF from 2001-2007. During that period, trawling effort declined 76.5 %: average annual reported trawling trips declined from 5,546 trips/year in 2001 to 1,303 trips/year in 2007, a decline in relative fishing effort from 4.26 to 1.0 (Figure 7). After calibration, we ran an EcoSim scenario beginning in 2001, incorporating the historical shrimp trawl fishery effort and catches reported to NCDMF from 2001-2007, and simulated a complete shrimp trawl ban (0% trawling effort) in 2008, ending in 2026 (a 25-year run). A trawl ban in estuarine waters has been recently proposed in North Carolina, and is under consideration at the current time by the North Carolina Fishery Commission, so this simulation is timely and reflects what could happen if a trawl ban were enacted. Time series and predictions form the Ecosim simulations are shown for some key fishery species (Figure 7) that are the most valuable fishery species in North Carolina ($13.3 million for pink, brown, and white shrimp combined, $21.8 million for blue crabs, $4.5 million for southern flounder (*Paralichthys lethostigma*) in 2012, NCDMF commercial landings data (40).

**Figure 7.**
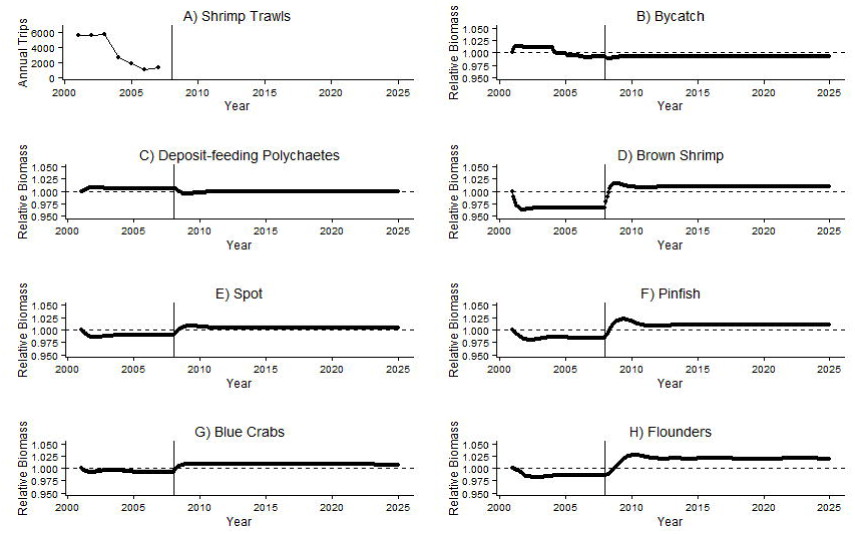
Ecosim projections of relative biomass (2001 baseline gC m^−2^), while simulating the impact of a trawl net ban (after 2008), for the bycatch and key species in Core Sound, NC. Top row, left: trawling trips reported to NCDMF; top row, right: bycatch (discards of all species); second row, left: benthic deposit-feeding polychaetes; second row, right: brown shrimp; third row, left: spot, *Leiostomus xanthurus*; third row, right: pinfish, *Lagodon rhomboides*; bottom left: blue crabs *Callinectes sapidus*, bottom right: southern flounder *Paralichthys lethostigma* and other paralichthid flounders. Dashed line represents the steady-state biomass in 2001 prior to the trawl ban for each compartment.

Consistent with our hypothesized trawling-induced trophic cascade, the closure of the shrimp trawling areas is predicted to cause some species to increase (spot and pinfish) and others to decline (deposit-feeding polychaetes and Penaeidae shrimp) after the simulated trawling ban. There was an increase (0.7175 g C m^−2^ biomass) in deposit-feeding polychaetes during year 2003 when trawling was at its peak in the historical time series. Our simulation model shows that the biomass of this scavenger group would decline after trawling cessation, reaching background relative biomass (0.7099 gC m^−2^) in 2009, a decline of 1.3% in biomass (Figure 7). This predicted decline after the simulated trawling ban is concurrent with a shrimp trawling effort decline and the increase of brown shrimp, other penaeid shrimp, and the bycatch species (primarily spot *Leiostomus xanthurus* and pinfish *Lagodon rhomboides* in Figure 7), all of which feed on polychaetes. These dramatic changes to the benthos and the fishes that feed on the benthos is due to the cessation of a trawling-induced trophic cascade, which depleted brown shrimp to 0.96 relative biomass during the peak of trawling effort (2001-2002). Thus, our simulation model appears to predict the direction of change in the standing stock of deposit-feeding polychaetes biomass, although not the magnitude of the difference in our measurements taken in open and closed trawling areas (Table 1).

It is notable in the trawl ban scenario on key fisheries in North Carolina: southern flounders and others in the family Paralichthyidae, after declining to 0.0256 gC m^−2^ due to trawling impacts in year 3 (2004), are predicted to return to the pre-trawling biomass of 0.0265 gC m^−2^ by year 6, due to declining fishing effort, then reach a peak biomass 15 years after a trawl ban, with only a 2% increase relative to 2001. This suggests that trawling ban would increase slightly the southern flounder stocks; this is a desirable effect, as the flounder stock is currently depleted. Spot (0.5%) and pinfish (0.9%) and brown shrimp (0.8%) would also increase slightly over 25 years. Finally, a trawl ban would actually cause decrease in the stocks of brown, pink and white shrimps, perhaps because of the increase in predation by other predators (flounders). There would be negligible increase of blue crabs (0.9% increase relative to 2001 biomass) followed by a slight decline (0.6673 gC m^−2^). The largest increase in biomass occurred in flounders, which are at the highest trophic levels of these key fishery species. Thus, a shrimp trawling ban is predicted to reverse the trophic cascade allowing biomass to pass back up the food web.

Fisheries managers in North Carolina are faced with a difficult choice based on these interacting fisheries: a trawl ban may cause a relatively large increase in in the biomass of the shrimp stocks, but this would be transfer energy to higher trophic level species, resulting in a slight increase the blue crab and larger increase southern flounder stocks. Shrimp would no longer be harvested by the trawlers, and any increase in their stocks would simply feed high trophic level fishes like the southern flounder, with the system reaching an equilibrium after 2018, a recovery period of 10 years. The fisheries for blue crabs, flounder, and shrimp are thus interconnected, directly by being harvested in the same gear (trawls) and indirectly through the food web network in this ecosystem, and cannot be easily managed separately. Note that this is true for the other fisheries (haul seines, gill nets, pound nets, crab pots) as well, which are still included in our ecosystem network models, but here modeled as having unchanging effort. These other fisheries may show increased fishing effort after a trawl ban, and increased harvests of these species, because fishermen will be likely to switch to the alternative gears (e.g., use more gill nets to catch increased flounder stocks) to target the same species, but this is not a scenario we have modeled.

## Discussion

Contrary to previous studies (cite them), there was more benthic scavenger biomass (deposit-feeding polychaetes) in the areas open to trawling after the shrimp season ended in the estuarine ecosystem in Core Sound. Further, in contrast to the earlier studies measuring acute trawling impacts on the benthos, the areas measured in the current study have been subjected to trawling for over thirty years. These findings are consistent with a long-term trawling-induced indirect effect or trophic cascade, which has been observed due to fishing in other ecosystems elsewhere (10–13,41) and as revealed by simulations in our ecosystem models. In a trophic cascade, the removal of a high-trophic-level species causes an increase in its prey species, which then decreases the abundance of that species’ prey. Shrimp trawling is a high-trophic level fishery (ETL=3.87)(21) that, as we observed, has reduced the abundance of benthic-feeding fishes (bycatch in the fishery is mostly pinfish and spot), and we suggest that their prey (deposit-feeding polychaetes) have increased as an indirect effect in the areas open to trawling, due to reduced fish predation, as a result of a trophic cascade.

What is unique about the trophic cascade in our system is that we now have a confirmation of a time-series based simulation model that mimics in a qualitative way the measurements of the prey’s difference in biomass between benthic samples taken in a marine protected area (closed trawling area) and a fished area (open trawling area). This system has been repeatedly trawled over thirty years and the discarded bycatch has been returned to the system (rather than exported to markets) and subsidized the detrital food web and benthos. This bycatch subsidy has resulted in an even greater difference in biomass of deposit-feeding polychaetes. It is important to note that other factors, such as water conditions in the closed areas (pollution for land run-off) and lack of mixing of the sediments by trawlers (20) after the no trawling areas were established in the 1970’s, could have resulted in environmental conditions that may contributed to of the difference in deposit-feeding polychaete biomass.

This trawling-induced trophic cascade hypothesis requires further experimental testing. The increase in benthos that has resulted directly from the trawling-induced trophic cascade influenced the whole ecosystem that produces the brown, pink and white shrimp of Core Sound because 1) the shrimp trawlers remove and return to the sea much of the juvenile fish biomass as dead discards, and harvest of penaeid shrimps and blue crabs reduces the overall predation on the benthos; and 2) the discard of the bycatch fishes as non-living carrion feeds the scavenger guild of Core Sound. The bycatch is fed upon by decomposing bacteria, microbes, polychaetes, penaeid and other shrimp, snails, and blue crabs and other crabs in these models. Indeed, other work suggests that δN^15^ is enriched the primary bycatch species pinfish and spot in the open trawling areas of Core Sound (25). In addition, this same study revealed an increase in the effective trophic levels for these bycatch species in the open trawling area Ecopath models (21). Our measurements of greater biomass of deposit-feeding polychaetes and lowered biomass of bycatch species (pinfish and spot), which were taken after the trawling season, in conjunction with the Ecopath/Ecosim simulation results, suggest that a trawling-induced trophic cascade has occurred in the study area. The observation that pinfish and spot in the open trawling areas were enriched in δN^15^ ratios after the trawling season suggests that a trophic subsidy occurred as well, due to carrion returned to the system, which added a partial increase in trophic level. The combined effect of the trophic cascade due trawling and the increase in biomass of the deposit-feeding polychaetes suggests a pronounced effect of shrimping on the whole ecosystem. Importantly, these measured shrimp-trawling impacts provide empirical verification of the Ecosim simulation model, which has not been accomplished previously.

There is a clear effect of shrimp trawling on the Core Sound ecosystem, causing a trawling-induced trophic cascade. The question remains if this is detrimental to the ecosystems’ functioning, and if societal goals for beneficial uses of Core Sound are being met. Based on the general perceived negative impact of trawling discards on these ecosystems, both in the USA and in Europe, North Carolina is considering enacting a shrimp trawl ban in estuaries. Our results call into question the general negative impact of trawl discards. Discards may in fact benefit particular trophic groups in the benthos, subsidizing their growth and production of polychaetes. This trawling-induced trophic cascade and potential bycatch subsidy to the benthos is eventually returned to the ecosystem as a pulse of carbon, stimulating energy and productivity of the benthos, which cascades up the food web and results in more spot, pinfish and southern flounder, albeit after many years, after the trawling virtually ceased in our Ecosim simulations.

Whether the nursery areas should be expanded to completely close trawling in the estuary remains a significant management concern. We could use our existing models to explore trawling policy options, simulating an expansion of the nursery areas using the Ecospace module in Ecopath, and following the approach of (42); however, this would require better spatially referenced harvest data for Core Sound. We recommend that fishery managers should proceed with caution and conduct an experiment, perhaps they should close some currently open trawling areas, and open some of currently closed the nursery areas to trawling, to see if this trophic cascade and bycatch stimulus effect can be measured. This should be considered as a temporary, and experimental, management option, with proper experimental protocols established to monitor the plankton, benthos, seagrasses, fishes, and larger vertebrates in an experimental design. We would recommend that long-term trawling closure-opening experiments be conducted with cooperation of the fishing industry, so the effects of shrimp trawling can be directly observed with rigorous before and after control and impact (BACI) study.

## Acknowledgements

The Coastal Resources Management Doctoral Program, the Institute for Coastal Science and Policy and the Department of Biology provided support for two of the authors (R. Deehr and K. Hart) financially while they were students at East Carolina University. We obtained support to conduct the field research and data analysis from North Carolina Sea Grant, project R/BS-1. We also thank Kyle Regensburg (Department of Biology at East Carolina University) for laboratory processing of benthic meiofauna samples; Cecilia Krahforst (Department of Biology at East Carolina University) for field collection help. Jill Luczkovich was a bird observer during our field studies. Robert Christian (Department of Biology at East Carolina University) reviewed an earlier draft of the manuscript. Finally, we are indebted to Carl Walters (University for British Columbia) helped us with Ecosim modelling advice.

